# New strategy to select cross-reactivity Meningococci strains: immunization with Outer membrane vesicles of serogroup C and cationic lipid as adjuvant

**DOI:** 10.1101/2021.05.17.444492

**Authors:** Amanda Izeli Portilho, Gabriela Trzewikowski de Lima, Elizabeth De Gaspari

## Abstract

**Background:** Invasive meningococcal disease (IMD), caused by *Neisseria meningitidis*, is a public health problem, associated with high levels of morbidity and mortality, capable of causing outbreaks or epidemics, but preventable through vaccination. In Brazil, the main serogroups isolated are C and B. The last epidemic occurred in the ‘80s, in São Paulo, because of a B:4:P1.15 strain.

**Methods:** Adult Swiss mice were immunized with outer membrane vesicles (OMV) of *N. meningitidis* strain C:4:P1.15, adjuvanted by the cationic lipid dioctadecyldimethylammonium bromide in bilayer fragments (DDA-BF), administered via prime-booster (intranasal/subcutaneous) scheme. The humoral response was accessed by Immunoblotting and ELISA, using homologous immunization strain and a different serogroup but equal serosubtype strain, *N. meningitidis* B:4:P1.15.

**Results:** Immunoblotting revealed the recognition of antigens associated with the molecular weight of Porin A and Opacity proteins, which are immunogenic but highly heterogeneous, and Tbp and NspA, which are more homogeneous between meningococci strains. ELISA results showed antibody production that persisted after 190 days and recognized the C:4:P1.15 and the B:4:P1.15 strains, with high avidity index. The adjuvanted group recognized antigens following the IN prime and had a higher avidity index against the heterologous strain.

**Conclusions:** DDA-BF improved the humoral response, but the OMV alone induced high avidity index antibodies as well. Even though these are preliminary results, we see it as a promising approach for affordable meningococcal immunization in developing countries, at outbreak or epidemic situations.

## Introduction

*Neisseria meningitidis* is the etiological agent of invasive meningococcal disease (IMD), which is associated with high morbidity and mortality rates [1]. The capsular polysaccharide and the content of the outer membrane vesicles (OMV) are important to trigger the immune response against the pathogen [2]. The immunization against serogroups A, C, W and Y is based on polysaccharide-protein conjugated vaccines, while serogroup B vaccination uses protein-based vaccines, given its poorly immunogenic polysaccharide, which could also lead to autoimmunity [3].

Mucosal immunization is promising in the prevention of infectious diseases by pathogens that firstly colonize mucous sites, as *N. meningitidis*, given that it promotes local and systemic immunity [4]. To promote adequate immune response after mucosal immunization, it is interesting to use adjuvants that contribute to the delivery and capture of the antigen [5, 6] The dioctadecyldimethylammonium bromide is a cationic lipid that has adjuvant characteristics and it is suitable to interact against the negative charged OMV of *Neisseriaceae* [7]. In this study, it was used in bilayer fragments (DDA-BF), that extend the surface contact between the adjuvant and the antigen and presents better stability [8].

It is important to observe the expression of immunogenic antigens when choosing a strain for vaccine production. The main limitation described for OMV vaccines is the strain-specific protection, related to the major expression of certain antigens [9]. However, outbreaks and epidemics are often caused by specific meningococci clones that presents little heterogeneity and can be controlled by OMV vaccines [10, 11]. It is interesting for countries that consider their adoption to keep epidemiological surveillance and phenotypical characterization of circulating meningococci strains, to maintain the conformity between the antigens expressed by the strains and the vaccine antigens [12]. In Brazil, the main isolates of *N. meningitidis* belong to serogroups C and B and the last epidemic occurred in the ‘80s, in São Paulo, because of a B:4:P1.15 strain [13]. Here, we tested the humoral immune response after immunization of Swiss mice with OMV of a Brazilian *N. meningitidis* C:4:P1.15 strain complexed with DDA-BF, through a prime-booster (intranasal/subcutaneous) immunization. It was used, as antigens, the homologous strain from immunization and a B:4:P1.15 strain, which presents a different serogroup but the same serotype and serosubtype and is representative of the last epidemic period and remained isolated from clinical samples until 2017 [13, 14].

## Methods

### *Neisseria meningitidis* strains and antigenic preparations

The *N. meningitidis* strain C:4:P1.15 was isolated from a Brazilian patient at the Bacteriology Center of Adolfo Lutz Institute (São Paulo, SP, Brazil), the Brazilian national reference laboratory for meningitis. To obtain the antigenic preparations, the strains were cultivated in Tryptic Soy Broth (TSB) (Difco Laboratories) supplemented with 1% of horse serum heat-inactivated (Sigma-Aldrich) for 24 hours at 37°C and 5%CO2 atmosphere [15]. For whole-cell suspensions of C:4:P1.15 and B:4:P1.15 strains, the bacterium was suspended, after cultivated, in Phosphate buffered-saline (PBS) containing 0.02% sodium azide and Optical Density (OD) were set for 0.1 at 620nm. For OMV of C:4:P1.15 strain, bacteria were incubated with a solution of 0.1M sodium acetate and 0.2M lithium chloride (pH 5.8) 2mm glass beads, than, taken to a shaker at 45°C, for 2 hours at 300 rpm. Afterwards, it was centrifuged at 8,000 rpm for 15 minutes at 4°C (Sorvall Instruments). The supernatant was detoxified from LPS using a Polymyxin B Sepharose 4B column [15].

The adjuvant DDA (Sigma-Aldrich) was diluted to 2mM, using 1mM saline solution. To obtain DDA on bilayer fragments (DDA-BF), it was performed sonication in 25% amplitude, for 15 minutes, at approximately 70°C (Sonics Vibrates Cell). Afterwards, the preparation was centrifuged for 30 minutes, at 10.000 rpm and 4°C (Eppendorf 5804R), for the removal of residual titanium [16].

Antigenic preparations were prepared with final concentrations: OMV 50µg/mL+DDA-BF 0.1mM and OMV 50µg/mL. Control groups were immunized only with DDA-BF 0.1mM, pre-immune sera were also used as control. OMV+DDA-BF complexes were characterized in size (374±67,7nm), polydispersion index (0,432±0,026) and zeta-potential (8,31±0.55) using ZetaPlus-Zeta Potential Analyzer (Brookhaven Instruments Corporation, Holtsville, NY) as described before [16].

### Immunization and blood collection

Adult females (4 months old) of Swiss mice were obtained from the bioterium facilities of Adolfo Lutz Institute (São Paulo, SP, Brazil) and kept in an experimental bioterium in the same location. All animal experimentation was performed following current legislation and was approved by the Ethics Committee on Animal Use of this institution (CEUA/IAL number 03/2012) and by the Technical and Scientific Committee of this institution (CTC/IAL number 4D-2011).

The mice received 4 intranasal (IN) doses (5µL each, 0.25µg of OMV) on days 1, 2, 22 and 23, and one subcutaneous (SC) booster dose (50µL, 2.5µg of OMV) on day 41, as an alternative to the classic prime booster schedule [17]. Blood was collected from the ophthalmic plexus before immunization (pre-immune bled), 20 days (prime bled) after IN doses 10 and 190 days (early and late booster bled, respectively) after SC dose, as described in Figure 1.

**Figure 1.**
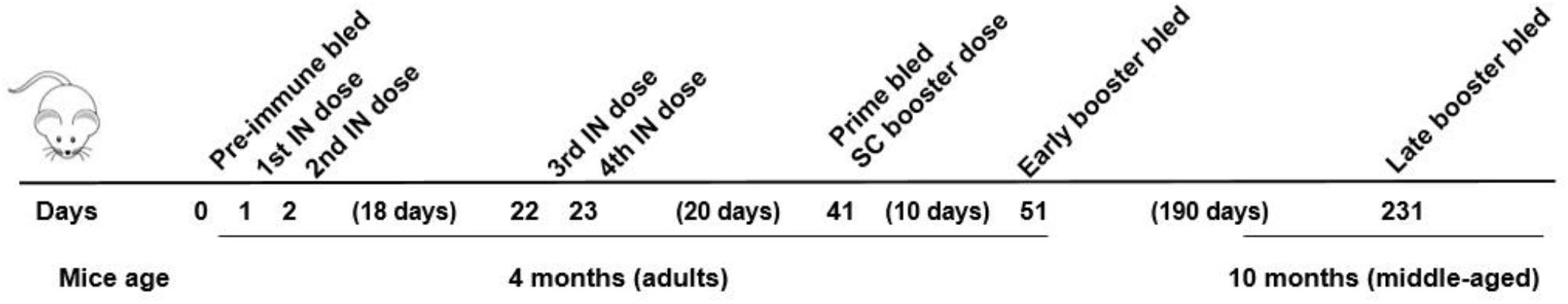
Immunization schedule and blood collection. (IN – intranasal; SC – subcutaneous)

### SDS-PAGE

The OMV was characterized by its electrophoretic profile in a 13% polyacrylamide gel in the presence of sodium dodecyl sulfate (SDS), along with a molecular weight (MW) marker (TrueColor High Range Protein Marker, Sinapse) ranging from 11 to 245 kDa. After the electrophoresis, the gel was stained with Coomassie Blue (PhastGel Blue R, Pharmacia Biotech)[18].

### Immunoblotting

Electrophoresis in 13% polyacrylamide gel was conducted using whole cells suspensions of C:4:P1.15 and B:4:P1.15 strains (approximately 1.5×109 cells each), as described above. After the electrophoresis, the protein was transferred to a nitrocellulose membrane of 0.45 µm (BioRad Laboratories) at 100 V, for 18 hours, at 4°C. Strips were blocked with Skim milk (La Sereníssima) 5%, overnight, at 4°C, and washed 5 times with PBS pH 7.2. Pooled sera (diluted at 1:25 in 2.5% Skim milk) obtained before the immunization, 20 days after IN dose, 10 days and 190 after SC dose,, was incubated overnight at 4°C, after, strips were washed 5 times with PBS. Anti-mouse IgG Fc-Horseradish-peroxidase labelled (HRP) (Kirkegaard & Perry Laboratories) diluted at 1:10.000 in Skim Milk 2.5% was incubated for 2 hours, at room temperature (∼25°C), under agitation. After 5 washes, it was added the substrate 3-Amino-9-ethylcarbazole (AEC) (Sigma-Aldrich), for 20 minutes, at room temperature, under agitation. The reaction was stopped by washing the strips with distilled water [11].

### ELISA

MaxiSorp ELISA plates (Nunc) were coated with whole-cell suspensions at OD 0.1 of *N. meningitidis* C:4:P1.15 strain or *N. meningitidis* B:4:P1.15 strain, and incubated overnight, at 37°C. The plates were washed 5 times with PBS pH 7.2-0.5% Tween 20, then, blocked with PBS-Skim Milk 5% (La Sereníssima) for 2h at 37°C. Individual mice serum collected 190 days after SC dose (diluted at 1:50 in PBS-Skim Milk 2.5%) was incubated for 2 hours at 37°C. After 5 washes, anti-mouse IgG Fc-Horseradish-peroxidase labelled (HRP) (Kirkegaard & Perry Laboratories) diluted at 1:20.000 in PBS-Skim Milk 2.5% was incubated for 2 hours at 37°C. The reaction was revealed by adding Tetramethylbenzidine (TMB) (Sigma-Aldrich) at 37°C for 20 minutes and stopped by the addition of sulfuric acid (H_2_SO_4_) 1N. OD was read at 450nm using a microplate reader (Multiskan Labsystems) [19].

### Avidity Index

To verify the avidity index (AI) of antibodies, it was performed a modified ELISA [20]. ELISA protocol was performed as described above, but it was added the chaotropic agent potassium thiocyanate (KSCN) (Sigma-Aldrich) at 1.5M, for 20 minutes, at room temperature (∼25°C), after the serum incubation and before the secondary antibody incubation. The AI was established as the ratio between the OD with KSCN and the OD without KSCN, multiplied by 100. It was classified as low if <30%, intermediate if between 30%-50% or high if >50% [21].

### Statistical analysis

Statistical analysis was performed using Kruskal-Wallis and Dunn’s post-test (GraphPad Prism 5, GraphPad Softwares). P<0.05 was considered statistically significant.

## Results

### SDS-PAGE and Immunoblotting

Figure 2a shows the electrophoretic profile of the OMV (C:4:P1.15) used for immunization. Figure 2b shows the Immunoblotting performed using the homologous and heterologous strains. Pre-immune sera were used as control. The group OMV+DDA-BF could recognize antigens following the IN prime, while the group OMV needed the SC booster for it. Given that the antigens of *N. meningitidis* are well described in the literature, it is possible to suggest, by the MW, which antigens were recognized. The recognized antigens of C:4:P1.15 strain presented around 100, 46, <32 and 22 kDa, which are related to Transferrin-binding protein (Tbp), Porin A (PorA), Opacity protein (Opa) and Neisserial surface protein A (NspA) [2]. In the case of B:4:P1.15 strain, only the antigen with <32 kDa was recognized.

**Figure 2.**
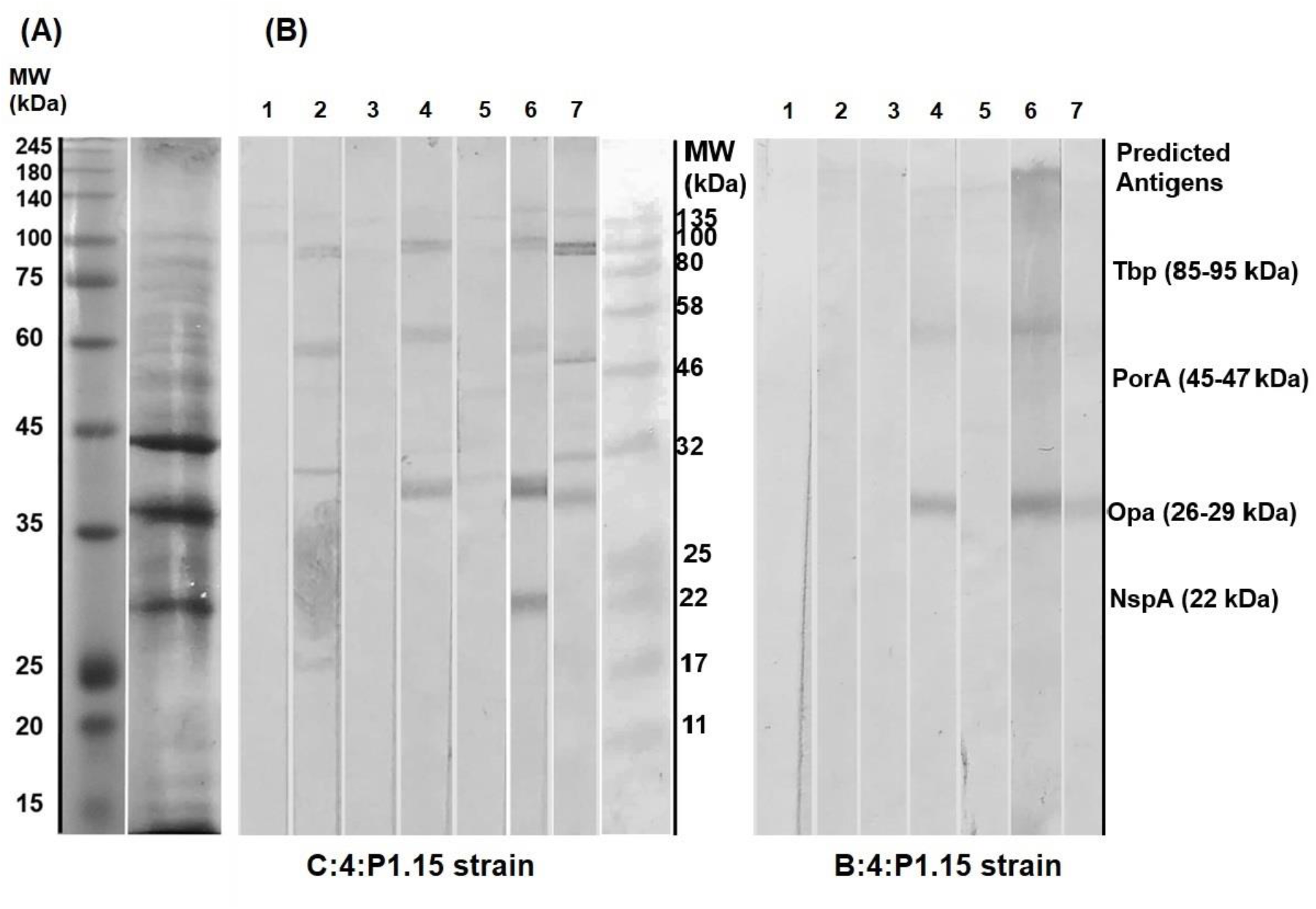
(A) Electrophoretic profile of OMV of strain C:4:P1.15, used for immunization. (B) Immunoblotting against homologous (C:4:P1.15) and heterologous (B:4:P1.15) strains, with 1. pooled pre-immune sera, 2. OMV+DDA-BF and 3. OMV pooled prime sera, 4. OMV+DDA-BF and 5. OMV pooled early booster sera, 6. OMV+DDA-BF and 7. OMV pooled late booster sera. (MW – Molecular Weight; kDa – kilodalton; Tbp – Transferrin-binding protein; PorA – Porin A; Opa – Opacity protein; NspA – Neisserial surface protein A). Antigens are only predicted by their MW and literature descriptions.

### ELISA and Avidity Index

Antibodies were observed against the C:4:P1.15 and the B:4:P1.15 strains (Figure 3a and 3b, respectively). Pre-immune and DDA-BF control presented lower OD. Only the group OMV+DDA-BF presented statically higher OD compared with pre-immune control, against both strains (p<0.05).

**Figure 3.**
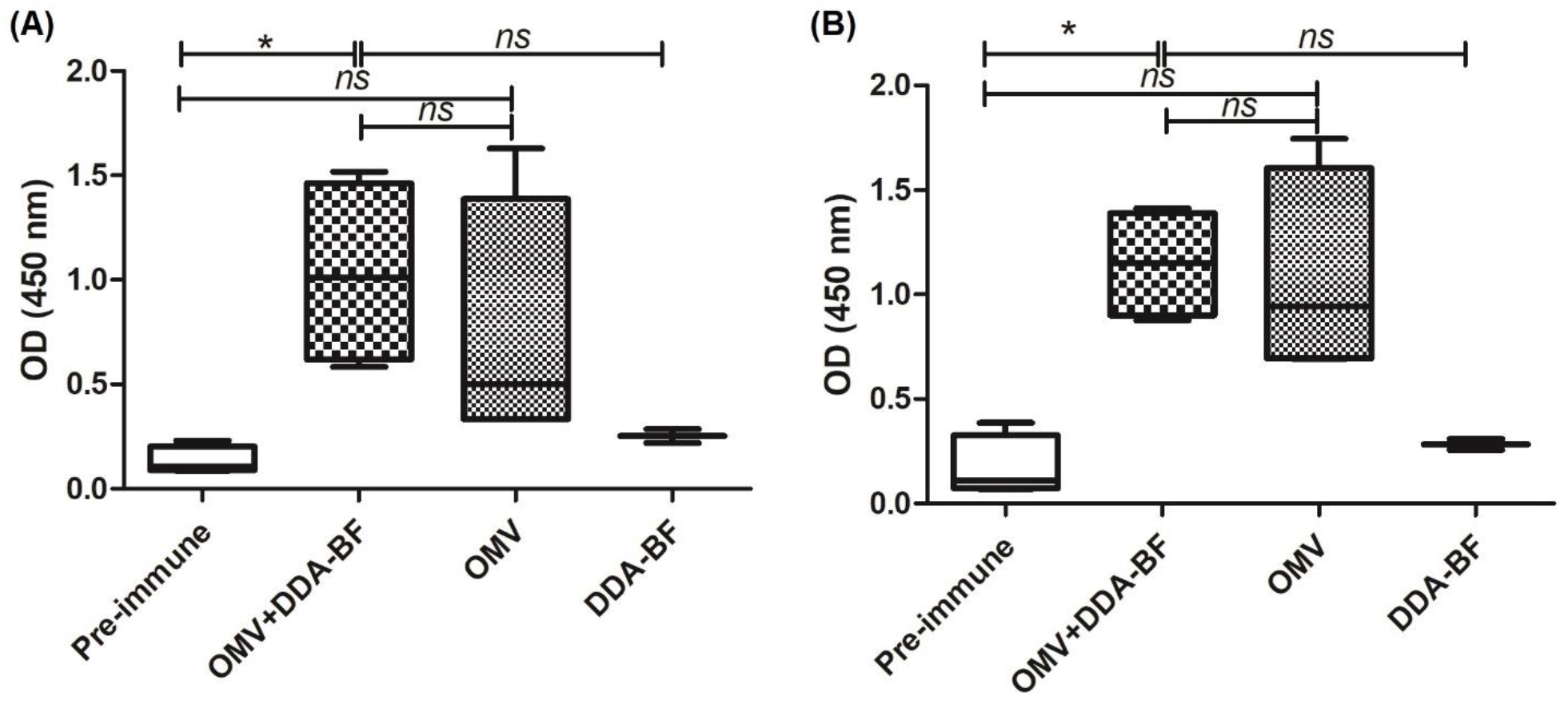
IgG levels in late booster sera of mice, diluted at 1:50, against (A) homologous strain (C:4:P1.15) and (B) heterologous strain (B:4:P1.15). (OD – Optical density; ns – not significant; *: p<0.05).

Table 1 presents the individual values for AI and the mean value of each group. OMV+DDA-BF and OMV groups presented high AI against both strains. The individual value varied between mice, however, all of them were considered high.

**Table 1.**
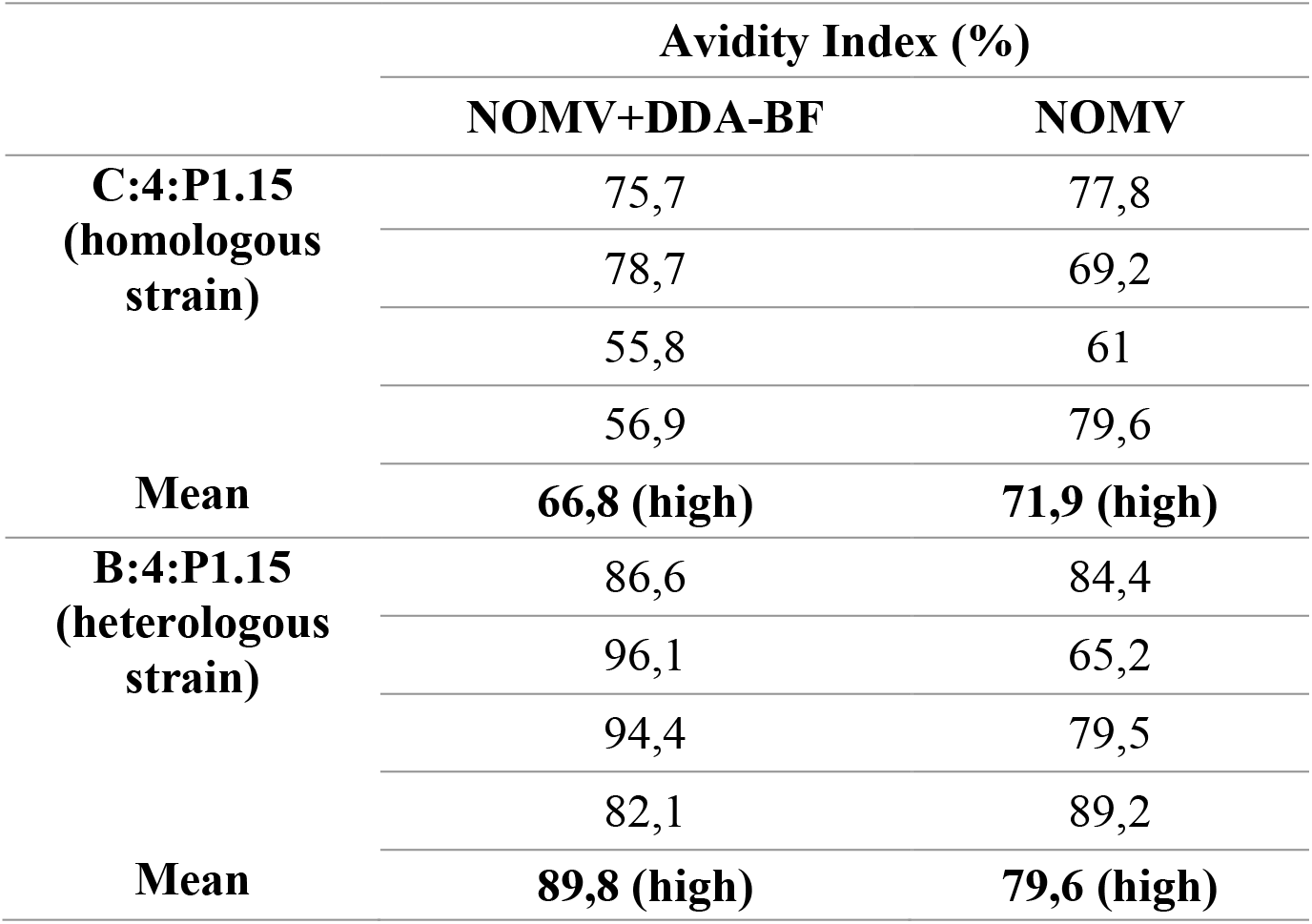
Individual and mean group value of AI against homologous and heterologous strain.

|  | Avidity Index (%) |  |
| --- | --- | --- |
|  | NOMV+DDA-BF | NOMV |
| **C:4:P1.15**<br>(homologous strain) | 75,7 | 77,8 |
|  | 78,7 | 69,2 |
|  | 55,8 | 61 |
|  | 56,9 | 79,6 |
| **Mean** | **66,8 (high)** | **71,9 (high)** |
| **B:4:P1.15**<br>(heterologous strain) | 86,6 | 84,4 |
|  | 96,1 | 65,2 |
|  | 94,4 | 79,5 |
|  | 82,1 | 89,2 |
| **Mean** | **89,8 (high)** | **79,6 (high)** |

## Discussion

The IN immunization offers mucosal and systemic immunity and mimics the course of natural infection, which could control contagious diseases from the site of infection [4]. The immunization schedule described here was designed to modify the prime booster scheme expecting that the interval between IN doses and the depot effect promoted by DDA-BF would ensure constant stimulus of the immune system [17, 22]. Another study of our group that used the prime-booster IN/SC scheme and DDA-BF as an adjuvant, but without interval between IN doses, verified a satisfactory humoral response [16]. Despite that, a study that directly compares the schedules, with and without interval between IN doses, using the same antigenic preparations, should be enrolled to confirm if one of them is better than the other.

The use of OMV in vaccines is well established against the serogroup B of *N. meningitidis*, since its capsular polysaccharide is poorly immunogenic and could induce autoimmunity [9]. OMV vaccines were used to control IMD outbreaks in Brazil, Cuba and New Zealand; the recombinant vaccine Bexsero (Glaxo Smith Kline) has been licensed in several countries and associates recombinant proteins with OMV from an epidemic *N. meningitidis* strain [23, 24]. The protein nature of OMV makes them an interesting antigen to achieve efficient, long-term immunity, given that it provides a T-dependent immune response [2]. The main limitation described for OMV vaccines is the immunodominance of Porin proteins on their content, which limits their cross-reactivity potential with strains that share the same serosubtype [9].

The immunization with OMVs from a C:4:P1.15 Brazilian strain, in this study, stimulated the production of antibodies against the same serosubtype but different serogroup strain B:4:P1.15, which is representative of a virulent clone that was responsible for the last meningococcal epidemic in São Paulo, Brazil, and was frequently isolated until 2017 [13, 14]. The homology between the porins of both strains probably ensured that antibodies trigged by one could recognize the other; however, different antigens might play a role in cross-reactivity. It is described that certain antigens, which are important for virulence, tend to be maintained, since it is observed that specific clones with elevated virulence often cause outbreaks and epidemics, which could be controlled by OMV vaccines [10]. This type of antigenic preparation is also cost and technically accessible to obtain, making it an option for rapid-response immunization, which is needed in epidemics situations.

Apart from that, other antigens present on OMV might be playing a role in the immune response: the OMV vaccine MenNZB, used in New Zealand, probably induced cross-protection against *N. gonorrhoeae* [25] and the vaccine Bexsero, composed of recombinant proteins and OMV from an epidemic strain, induced antibodies that were able to kill meningococci strains from different serogroups [26].

DDA-BF is a cationic lipid, which enhances the interaction of this adjuvant with antigen-presenting cells (APCs) [8]. The adjuvant seemed to improve the immune response, given that in Immunoblotting with C:4:P1.15 strain, the group immunized with OMV+DDA-BF recognized antigens following the IN priming, while the group immunized with OMV alone needed the booster dose to recognize the antigens. The adjuvanted group was able to recognize antigens from the heterologous strain sooner than the OMV group as well. Previous studies of our group showed that booster doses and adjuvant use improve antigenic recognition [11].

Based on the literature descriptions, we believe that the antigens recognized referred to Tbp, PorA, Opa and NspA. It is important to point that specific tools, like Monoclonal antibodies or molecular characterization, should be employed to confirm if these are the antigens present on Immunoblotting strips [11]. It was expected that the sera would recognize an antigen ∼46 kDa on both homologous and heterologous strain, because it is the MW related to PorA, that usually dominates the OMV content and is shared by the strains used. PorA antibodies present bactericidal capacity and confer protection [2], however, the antigen recognized in both strains is possibly Opa. This protein is responsible for adherence and invasion of host cells, and mediates immune interactions, being able to inhibit T and B lymphocyte activation. Anti-Opa antibodies are potentially bactericidal and block adhesion [27]. This protein proved to be immunogenic in recombinant vaccines [27]. Taken together, it led us to believe that the presence of such antigen would be interesting in the antigenic preparation and may confer a good immune response.

However, Opa presents an antigenic heterogeneity [15], which may be a limitation for broader coverage against strains, but can be helpful to control outbreaks. Tbp mediates the iron uptake from the host, which is needed for meningococcal survival, and presents limited heterogeneity [2]. Even in low titers, antibodies anti-Tbp were able to inhibit iron uptake and growth of homologous strain [28]. When used on immunization, these proteins presented promising results in animals, eliciting bactericidal antibodies against homologous and heterologous strains, which may be of interest for broader protection [29, 30]. NspA also presents limited heterogeneity, being interesting for cross-reactivity, and is capable of inducing bactericidal antibodies [2]. This protein, however, is a human Factor H ligand, which would impair its bactericidal capacity. Therefore, to confer protection, it should be associated with other antigens or manipulated to diminish its affinity by Factor H [31, 32].

Long-lasting immunity is another desired characteristic of vaccines [3]. The OMV+DDA-BF group was the only one to show a statistical difference when compared with pre-immune control (p<0.05), regardless of the strain used as coat antigen, restating the importance of adjuvants. It is interesting to point the sera used was collected 190 days after the booster dose, showing the persistence of immune response: mice were adult when immunized and middle-aged when sera were collected [33]. The persistence of immune response can be related to the immunogenicity of the OMV, but also to the presence of DDA-BF, which improve antigen presentation to APCs by its particulate form and slow liberation [8]. Especially in mucosal immunity, the use of adjuvants to ensure the protection, delivery and presentation of the antigen is an important subject [4]. It should be noted that despite the limited antigenic recognition in Immunoblotting, the OMV group showed a high OD in ELISA. It could be related to the presentation of the antigens on each assay: while Immunoblotting denatures the antigens [18], ELISA maintains the natural presentation of it, which could lead to better recognition by antibodies.

The avidity index has been described as a tool to infer bactericidal activity, however, classic serum bactericidal activity assay (SBA) should be enrolled to confirm the functionality of antibodies [34]. Given our results, present in table 1, it was interesting to observe that all animals presented a high AI against both strains. In our experience, OMV without adjuvants usually presents lower AI when compared with OMV complexed with adjuvants [7, 16, 35]. OMV can be highly immunogenic *per se*, but detoxification can decrease its immunogenicity, since lipopolysaccharide is one of the main immune activators [9]. In this study, OMV of the C:4:P1.15 strain alone induced less quantity of antibodies, but with high AI, suggesting that the strain maintained a high immunogenicity level despite the LPS detoxification, which makes it a promising strain for vaccine development. Even though, it should be pointed that it was a preliminary study and the strain and antigenic preparation were obtained in laboratory-based conditions. Sometimes, the antigenic presentation of a bacterial strain can change when it goes to large-scale production, decreasing the immunogenic potential [24]. In this scenario, the importance of adjuvants to improve immunogenicity is restated.

Despite its worldwide distribution, IMD presents a greater impact in low-income countries and immunization is the better strategy to control it, therefore, less expensive techniques to produce vaccines cannot be put aside [3]. One of the concerns around recombinant vaccines is their cost to be implemented in national programmes [36]. The technical and economic aspects to extract OMV are more accessible when compared to recombinant proteins. It is important to keep researching about them as cost-effective options for developing countries, as the efforts that had led to MenAfriVac and the current attempt to develop a low-cost quadrivalent vaccine against serogroups A, C, W and Y for use in Africa [37].

In Brazil, the epidemiology of IMD has been changing, with the increase of serogroup W in the South region, which accompanies the widespread of W:2a:P1.5,2 clone in South America and Europe [1,14]. Once again, it is important to maintain epidemiologic surveillance to follow these changes and adequate the immunization programs [1,12]. Our laboratory followed this strategy of cross-reactive immunization using OMV of C:2a:P1.5 strains against both C and W serogroups.

## Conclusion

According to our results, there is potential in this immunization approach, which induced persistent serum antibodies with high AI against Meningococci C and B strains expressing the same serosubtype. However, these are preliminary results. Assays that check the functionality of antibodies are needed to conclude if the immunization was protective. Since we observed some variation in immune response between individuals, it would be interesting to increase the experimental groups. The use of higher antigenic doses might also contribute to achieve a sustained response in all individuals. In epidemics caused by specific strains, the use of a non-manipulated strain that conserves its antigen’s expression and a scheme that provides mucosal and systemic response for an affordable immunization can be beneficial.

## Acknowledgements

The authors would like to thank Dr Ana Paula Silva Lemos, from Bacteriology Center of Adolfo Lutz Institute, for providing the bacteria strains used in this work and Dr Nilton Lincopan, from Institute of Biomedical Sciences of University of São Paulo for supporting DDA-BF preparation.

